# Population and comparative genetics of thermotolerance divergence between yeast species

**DOI:** 10.1101/2020.08.31.275453

**Authors:** Melanie B. Abrams, Claire A. Dubin, Faisal AlZaben, Juan Bravo, Pierre M. Joubert, Carly V. Weiss, Rachel B. Brem

## Abstract

Many familiar traits in the natural world—from lions’ manes to the longevity of bristlecone pine trees—arose in the distant past, and have long since fixed in their respective species. A key challenge in evolutionary genetics is to figure out how and why species-defining traits have come to be. We used the thermotolerance growth advantage of the yeast *Saccharomyces cerevisiae* over its sister species *Saccharomyces paradoxus* as a model for addressing these questions. Analyzing loci at which the *S. cerevisiae* allele promotes thermotolerance, we detected robust evidence for positive selection, including amino acid divergence between the species and conservation within *S. cerevisiae* populations. Since such signatures were particularly strong at the chromosome segregation gene *ESP1*, we used this locus as a case study for focused mechanistic follow-up. Experiments revealed that, in culture at high temperature, the *S. paradoxus ESP1* allele conferred a qualitative defect in biomass accumulation and cell division relative to the *S. cerevisiae* allele. Only genetic divergence in the *ESP1* coding region mattered phenotypically, with no functional impact detectable from the promoter. Together, these data support a model in which an ancient ancestor of *S. cerevisiae*, under selection to boost viability at high temperature, acquired amino acid variants at *ESP1* and many other loci, which have been constrained since then. Complex adaptations of this type hold promise as a paradigm for interspecies genetics, especially in deeply diverged traits that may have taken millions of years to evolve.

## INTRODUCTION

A central goal of research in evolutionary genetics is to understand how new traits are built. Much of the literature to date focuses on adaptive trait innovation within a species, in the wild (Asgari et al., 2020; Chan et al., 2010; Cleves et al., 2014; Field et al., 2016; Linnen et al., 2013; Will et al., 2010) and in the lab (Blount et al., 2012; Castro et al., 2019; Good et al., 2017; Tenaillon et al., 2016). These systems have enabled studies of short-term adaptation, its genetics (Barroso-Batista et al., 2014; Castro et al., 2019; Garud et al., 2015; Good et al., 2017; Harris et al., 2018; Xie et al., 2019) and its dynamics (Blount et al., 2012; Toprak et al., 2011). Such work on recent adaptations serves as a backdrop for the study of evolution over longer timescales. Many familiar traits from the natural world have been acquired over millions of generations. In the modern day, such characters manifest as differences between deeply diverged, reproductively isolated lineages. They can represent the abiding fitness strategies of their respective species, and are thus of particular interest in the field. But their evolutionary mechanisms pose a key challenge, given that the relevant events happened so long ago. For these ancient traits, candidate-gene studies have implicated individual loci (Anderson et al., 2016; Baldwin et al., 2014; Li and Fay, 2017; Liu et al., 2018; Massey and Wittkopp, 2016; Sackton et al., 2019; Sulak et al., 2016; Tian et al., 2019) and reconstructed the mutational path by which a given determinant evolved (Anderson et al., 2015; Bridgham et al., 2009; Finnigan et al., 2012; Liu et al., 2018; Pillai et al., 2020). Even in such landmark cases, the tempo and mode of evolution of deep trait divergences have remained largely out of reach. To meet the latter challenge, one would need to trace the rise of causal alleles in the respective species and the selective forces that drove it, and pinpoint the timing of these events.

In prior work, our group mapped multiple housekeeping genes underlying the difference in thermotolerance between *Saccharomyces cerevisiae* and other species in its clade, and found that *S. cerevisiae* harbored derived alleles at these loci (Weiss et al., 2018). Here we set out to investigate when and how *S. cerevisiae* acquired the putatively adaptive determinants of thermotolerance, using a population-genomic approach. We then used the results as a jumping-off point for additional analyses of the molecular mechanisms by which variants at thermotolerance genes confer their effects.

## MATERIALS AND METHODS

### Identifying thermotolerance genes

To identify genes at which variation between *S. cerevisiae* and *S. paradoxus* impacts thermotolerance, we re-analyzed data from a reciprocal hemizygosity screen of transposon mutants in the interspecies hybrid background (Weiss et al., 2018) as described, with the following differences. Call a_39,i_ the average, across technical replicates, of sequencing-based abundances of a hemizygote mutant measured after ∼7 generations in biological replicate *i* of growth at 39°C, and a_28,i_ the analogous quantity for growth at 28°C, for i = [1,3]. We calculated the mean of the latter across biological replicates, a_28,mean_, and then used it to tabulate three replicate estimates of the temperature effect on growth of the mutant as log_2_(a_39,i_/a_28,mean_). If the coefficient of variation across these biological replicates was greater than 20, we eliminated the mutant from further consideration. Otherwise, for a given gene, we concatenated these vectors of length three across all hemizygote mutants in the *S. paradoxus* allele for which we had abundance data, yielding the set of temperature effects s_Spar_. We did likewise for the *S. cerevisiae* allele, yielding s_Scer_. We retained for further analysis only genes at which we had at least two mutants’ worth of data for each allele. For each such gene, we compared s_Scer_ and s_Spar_ with a Wilcoxon test, and corrected for multiple testing across genes, as described in (Weiss et al., 2018).

### Sequence data, alignments, and interspecies diversity

For D_XY_ analyses in Table S2, for a given gene, open reading frame sequences for the strains of each *S. cerevisiae* population from (Peter et al., 2018) were aligned against the European *S. paradoxus* population from (Bergström et al., 2014) and, separately, against the North American *S. paradoxus* subpopulation B from (Durand et al., 2019). For D_XY_ analysis across species in Table 1, alignments were generated using all *S. cerevisiae* and *S. paradoxus* strains. Alignments used MUSCLE (Edgar, 2004) with the default settings for DNA and --maxiters set to 2. Any gene for which, in the alignment, >10% of sites were denoted as gaps or unknown nucleotides (Ns), or sequences from <75% of strains in the population were available, was eliminated from analysis, leaving 4110 to 4781 genes suitable for testing in each analysis.

**Table 1.**
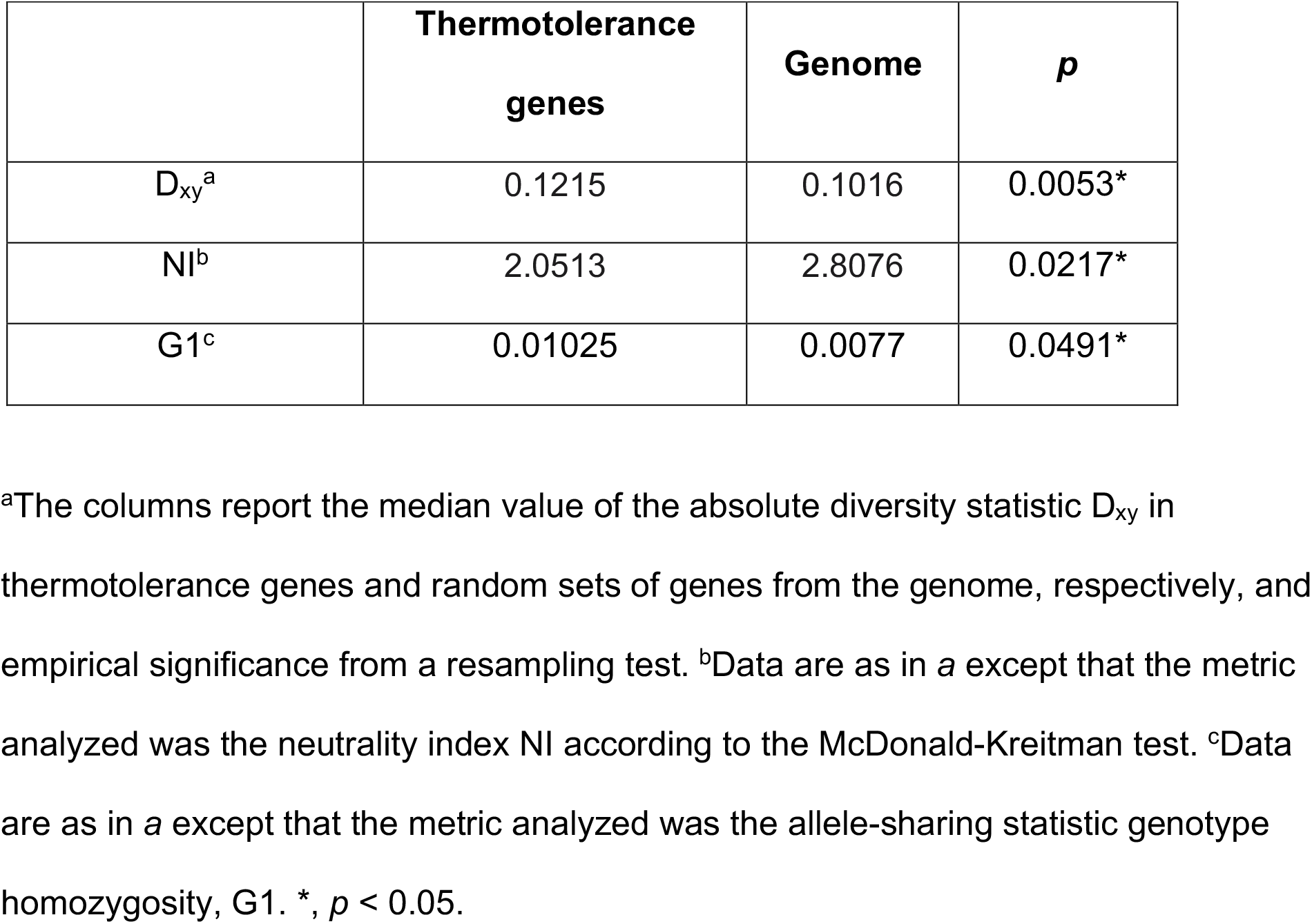
Thermotolerance loci are enriched for positive selection between species and allele-sharing in *S. cerevisiae*.

We calculated pairwise nucleotide diversity (D_xy_) for each gene as

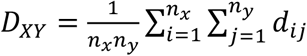

using a custom script, where *n*_*x*_ is the number of *S. cerevisia*e strains, *n*_*y*_ is the number of *S. paradoxus* strains, and *d* is the number of sites with nucleotides differences at the same position for each pairwise sequence comparison. Sites with gaps or unknown nucleotides were ignored.

To test for enriched D_xy_ among thermotolerance genes in a given *S. cerevisiae* and *S paradoxus* population pair or across the species, we first tabulated D_therm_, the median D_xy_ across the thermotolerance gene cohort from the appropriate genomes. We next sampled 10,000 random cohorts of genes from the genome with the same number of essential and nonessential genes as in the thermotolerance cohort (Winzeler et al., 1999), and tabulated the median D_xy_ in each D_rand_ from the appropriate genomes. We used as an empirical *p*-value the proportion of random cohorts with D_rand_ ≥ D_therm_.

### Codon alignment and McDonald-Kreitman statistics

Open reading frame sequences for each *S. cerevisiae* strain from (Peter et al., 2018), the European *S. paradoxus* strains (Bergström et al., 2014), and North American *S. paradoxus* strains (Durand et al., 2019) were translated to amino acid sequences using Biopython (Cock et al., 2009) and aligned using MUSCLE (Edgar, 2004) with default settings for amino acids and --maxiters set to 2. The amino acid sequence alignments and unaligned nucleotide sequences were used as input to PAL2NAL (Suyama et al., 2006) to create codon alignments for each gene. Sequences with stop codons within the open reading frame or where >10% of sites were denoted as gaps or unknown nucleotides (Ns) were discarded. Genes with valid sequences in <75% of strains from each species were removed from the analysis, leaving 3814 genes suitable for testing.

The codon alignments were input into the CodonAlignment module of Biopython 1.78 (Cock et al., 2009) and the mktest function reported the number of divergent nonsynonymous (*D*_*n*_), divergent synonymous (*D*_*s*_), polymorphic nonsynonymous (*P*_*n*_), and polymorphic synonymous changes (*P*_*s*_) in each gene. We calculated the Neutrality Index (NI) for each gene as

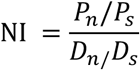

(McDonald and Kreitman, 1991). We then used these measures as input into a resampling test for enrichment of low NI, analogous to that used for D_xy_ (see above).

### Multi-locus genotype and allele-sharing inference in *S. cerevisiae* populations

We calculated expected genotype homozygosity, G1 (Harris et al., 2018), as follows. For the allele-sharing inference across all *S. cerevisiae* in Table 1, we used unphased VCF genotypes for all strains from 1011 Yeast Genomes (Peter et al., 2018) as input into SelectionHapStats (Harris et al., 2018) with the following parameters: -w (window size, SNPs) = 1200, and -j (jump size, SNPs) = 25. We tabulated G1 in each window whose center fell within the open reading frame, and we calculated the average across the windows. We then used these measures as input into a resampling test for enrichment of high G1 analogous to that used for D_xy_ (see above). For allele-sharing inference in individual populations of *S. cerevisiae* in Table S3, we proceeded as above except that we used unphased VCF genotypes for each of the five largest *S. cerevisiae* populations from 1011 Yeast Genomes (Peter et al., 2018).

We also evaluated allele-sharing at thermotolerance genes in *S. paradoxus*, as a complement to the above analyses in *S. cerevisiae*. For this purpose we used unphased VCF genotypes for all *S. paradoxus* genomes from (Bergström et al., 2014) according to the methods above and found no significant enrichment for G1 at thermotolerance loci (resampling *p* = 0.19).

### Polymorphism in Wine/European *S. cerevisiae*

For polymoprhism calculations in Figure S1, from the Wine/European *S. cerevisiae* population from (Peter et al., 2018), we used genotype data as a VCF as input into VCFtools (Danecek et al., 2011) with the command --site-pi. This output a polymorphism (π) value for each SNP. We tabulated the average π across a 1200 SNP window centered at each SNP, with invariant sites contributing π = 0.

### *ESP1* phylogenetic analysis

We used the alignment of the open reading frame of *ESP1* from the type strains of *S. cerevisiae, S. paradoxus, S. mikatae, S. bayanus, S. uvarum*, and *S. kudriavzevii* from saccharomycessensustricto.org as input into the codeml module of PAML4.9 (Yang, 2007). The branch-site model (model=2, NSsites=2) was used, and two models, null and alternative, were fitted. In the null model, the dN/dS for the *S. cerevisiae* branch was fixed at 1.0 and all other branches were described by the same dN/dS ratio (ω). In the alternative model, the branch leading to *S. cerevisiae* was fitted with one ω, and all other branches were fitted with a separate ω. A test statistic, calculated by comparing the likelihood ratios of the alternative and null models, was used to calculate a *p*-value by comparing it to a chi-squared distribution with one degree of freedom, equal to the difference in the number of parameters in the two models. No codons exhibited a posterior probability of positive selection, on the branch leading to *S. cerevisiae*, higher than 0.9.

### Analysis of *cis*-regulatory expression divergence between *S. cerevisiae* and *S. paradoxus*

For Table S5, we analyzed temperature-dependent allele-specific expression measurements in interspecific hybrids as follows. For each gene in turn, from (Tirosh et al., 2009) we tabulated the log_2_-ratio of allele-specific expression between alleles of an *S. cerevisiae* x *S. paradoxus* hybrid cultured at 35°C, as a difference from the analogous quantity from cultures at 30°C, which we refer to as D_ASE_. To test for an enrichment of high-magnitude allele-specific expression differences between species at thermotolerance loci, we took the absolute value of D_ASE_ for each gene and then tabulated the median of this value across the thermotolerance gene set, D_ASE,therm_. We next sampled 10,000 random cohorts of genes from the genome with the same number of essential and nonessential genes as in the thermotolerance cohort (Winzeler et al., 1999), and for each we calculated the median D_ASE_, D_ASE,rand_. We then used as an empirical *p*-value the proportion of random cohorts for which D_ASE,rand_ ≥ D_ASE,therm_. To test for directional *cis*-regulatory change between species at thermotolerance genes, we repeated the above analysis except that we took the median across signed D_ASE_ values for a gene set of interest, and then used as an empirical *p*-value the proportion of random cohorts at which |ASE_rand_| ≥ |ASE_therm_|.

Separately, we repeated the above analysis using measurements from 37°C and 33°C cultures of an *S. cerevisiae* x *S. uvarum* hybrid from (Li and Fay, 2017), which are reported as allele-specific expression a_T,s_ for the allele from species *s* at temperature *T*. We tabulated D_ASE_ for each gene as log_2_(a_37,Scer_/a_37,Suv_) - log_2_(a_33,Scer_/a_33,Suv_) and tested for enrichment of high-magnitude and directional *cis*-regulatory variation across thermotolerance genes as above.

### Interspecies swap strain construction at *ESP1* promoter and coding region

To swap the allele of the *ESP1* promoter from *S. paradoxus* Z1 into *S. cerevisiae* DBVPG1373, and likewise for the coding region, we designed allele-specific Cas9 guide RNAs for the *S. cerevisiae* background, generated donor DNA from *S. paradoxus*, transformed, and screened for successful transgenesis by Sanger sequencing as in (Weiss et al., 2018). Strains are listed in Table S6.

### Large-format growth assay

For growth measurements in Figures 2 and S2, we assayed *S. paradoxus* Z1, *S. cerevisiae* DBVPG1373, the full *ESP1* swap in the *S. cerevisiae* background (harboring the promoter and open reading frame from *S. paradoxus*) from (Weiss et al., 2018), and the *ESP1* promoter and coding swaps in the *S. cerevisiae* background (see above) as follows. Each strain was streaked from a -80°C freezer stock onto a yeast peptone dextrose (YPD) agar plate and incubated at room temperature for 3 days. For each biological replicate, a single colony was inoculated into 5 mL liquid YPD and grown for 24 hours at 28°C with shaking at 200 rpm to generate pre-cultures. Each pre-culture was back-diluted into YPD at an OD_600_ of 0.05 and grown for an additional 5.5-6 hours at 28°C, shaking at 200 rpm, until reaching logarithmic phase. Each pre-culture was again back-diluted into 10 mL YPD in 1-inch diameter glass tubes with a target OD_600_ of 0.05; the actual OD_600_ of each was measured, after which it was grown at either 28 or 39°C with shaking at 200rpm for 24 hours, and OD_600_ was measured again. The growth efficiency for each replicate was calculated as the difference between these final and initial OD_600_ values. The pipeline from inoculation off of solid plates through preculture, two back-dilutions, and growth at 28 or 39°C we refer to as a day’s growth experiment. For each day’s experiments, we calculated the average efficiency <e_Scer_> across the replicates of wild-type *S. cerevisiae*, and we used this quantity to normalize the efficiency e_s_ measured for each replicate assayed on that day of a genotype of interest *s*. Thus, the final measurement used for analysis for each replicate on a given day was e_s_/<e_Scer_>. We carried out a total of 2-3 days’ worth of replicate growth experiments for each genotype, with three separate transformant strains analyzed by this workflow in the case of the coding swap. For a given genotype we used the complete cohort of measurements of e_s_ /<e_Scer_> from all days and strains as input into a one-sample, one-tailed Wilcoxon test to evaluate whether e_s_/<e_Scer_> was less than 1 (*i*.*e*. that the strain grew worse at 39°C than wild-type *S. cerevisiae*).

### Temperature dose-response growth assay

To evaluate temperature dose-responses in Figure 3, we assayed *S. paradoxus* Z1, *S. cerevisiae* DBVPG1373, and the full *ESP1* swap in the *S. cerevisiae* background (harboring the promoter and open reading frame from *S. paradoxus*) from (Weiss et al., 2018) as follows. Each strain was streaked from a -80°C freezer stock onto a YPD agar plate and incubated at room temperature for 3 days. For each biological replicate, a single colony was inoculated into 5 mL liquid YPD and grown for 48 hours at 28°C with shaking at 200 rpm to create a stationary phase pre-culture. From pre-culture we made eight back-dilution experimental cultures in a standard PCR strip tube, each in 200 µL YPD, and we incubated these in a thermocycler using a gradient protocol from 37.0 to 40.8°C. After 24 hours, 150 µL from each culture was removed and OD_600_ was measured. The pipeline from inoculation off of solid plates through pre-culture, back-dilution, and growth we refer to as a day’s growth experiment for the dose-response of a strain. For each day’s experiments, at a given temperature we calculated the average efficiency <e_Scer,37_> across the replicates of wild-type *S. cerevisiae* at 37°C, and used it to normalize the efficiency e_s,T_ measured for each replicate assayed on that day of a strain of interest *s* at temperature T. Thus, the final measurement used for analysis for each replicate and temperature on a given day was e_s,T_/<e_Scer,37_>. We carried out two days’ worth of replicate growth experiments, and used the complete cohort of measurements of e_s,T_/<e_Scer,37_> from all days and all temperatures as input into a two-factor type 2 ANOVA test for a temperature-by-strain effect comparing *s* with *S. cerevisiae*.

### Microscopy

Microscopy was performed as described in (Weiss et al., 2018). Images were scored, blinded, for the size of dyads, omitting all clumps of >2 cells. Two replicates per strain and condition were imaged, and ten to sixteen images per replicate were scored. Significance was evaluated using a two-factor ANOVA test to evaluate strain by temperature effects. The range and mean number of dyads scored per image and per strain are reported in Table S7.

## DATA AVAILABILITY

Strains and plasmids are available upon request. The authors affirm that all data necessary for confirming the conclusions of the article are present within the article, figures, and tables. Custom scripts for sequence preparation and population genetics statistics for absolute sequence divergence, absolute nucleotide divergence, and the McDonald-Kreitman neutrality index are available at https://github.com/clairedubin/thermotolerance. Custom scripts for RH-seq reanalysis, multi-locus genotype and allele-sharing inference, and allele-specific expression analysis are available at https://github.com/melanieabrams-pub/thermotolerance-loci-across-yeasts.

## RESULTS

### Signatures of adaptation and constraint at *S. cerevisiae* thermotolerance loci

With the goal of investigating evolutionary mechanisms of thermotolerance divergence between yeasts, we started by addressing the genetics of the trait. Our earlier study used genome-scale screens with the reciprocal hemizygosity test (Steinmetz et al., 2002; Stern, 2014) to identify eight genes at which *S. cerevisiae* harbored pro-thermotolerance alleles relative to those of *S. paradoxus* (Weiss et al., 2018). We re-processed these screen data with an improved statistical workflow, to boost power and genome coverage (see Methods). The results recapitulated seven loci that we had reported and validated, plus an additional seven that had not risen to significance in our original analysis (Table S1). We considered the expanded set of loci as a more complete model of the genetic architecture of the trait, which would be well-suited to population and evolutionary analyses.

Thermotolerance is a defining and putatively adaptive character of *S. cerevisiae*, shared among isolates within the species and distinguishing it from the rest of the *Saccharomyces* clade (Gonçalves et al., 2011; Salvadó et al., 2011; Sweeney et al., 2004). We hypothesized that the loci underlying thermotolerance had evolved under positive selection before the radiation of modern *S. cerevisiae* populations. To test this, we made use of a broad population survey of *S. cerevisiae* (Peter et al., 2018), and the deepest-sampled *S. paradoxus* populations available (from vineyards and European collection locales (Bergström et al., 2014) and from North America (Durand et al., 2019)). With these genomes, we first sought to quantify sequence diversity between the species, at thermotolerance genes. The absolute diversity statistic D_xy_ reaches high levels in a lineage after selection when compared to a representative of the ancestral state (Nei, 1987), and is preferred over relative-divergence metrics as a suggestive statistic of adaptation (Noor and Bennett, 2009). Using the entire set of population genomes from *S. cerevisiae* and S. *paradoxus*, we found enrichment for high D_xy_ among our thermotolerance genes (Table 1 and Table S2), as expected from a previous smaller-scale analysis (Weiss et al., 2018). The latter result was mirrored by analyses of individual *S. cerevisiae* populations (Table S3), ruling out demographic artifacts as the source of signal in our species-wide test. Thus, divergence from *S. paradoxus* at thermotolerance loci is a trend that pervades >30 *S. cerevisiae* populations, collected in Europe, Asia, Africa, and the Americas, supporting a model of a selective event in the ancestor of modern *S. cerevisiae*.

We next reasoned that, if evolution had used predominantly amino acid variants in building the thermotolerance trait, the underlying loci would exhibit striking coding variation between species, relative to within-species polymorphism and relative to synonymous changes, as analyzed in the family of methods derived from the McDonald-Kreitman test (McDonald and Kreitman, 1991). For enrichment tests we used the neutrality index (Stoletzki and Eyre-Walker, 2011), which reaches low values in cases of adaptive amino-acid evolution between species. The results revealed a 1.37-fold reduced neutrality index among thermotolerance genes relative to the genome as a whole (Table 1 and Table S2). Such a signal strongly supports a history of adaptation at thermotolerance loci, with a mechanism involving changes to protein structure and/or function.

Under our model of thermotolerance evolution, after an ancestral *S. cerevisiae* population gained the trait long ago, it was maintained by purifying selection throughout the species. To assess signatures of constraint within *S. cerevisiae* on thermotolerance loci, we reasoned that haplotype-level analyses would have greater power than site-by-site tests. The SelectionHapStats suite (Garud et al., 2015; Harris et al., 2018) can be used for this purpose to detect very recent soft selective sweeps in young populations or, for a putatively ancient adaptation like ours, to report conservation more generally. In analyses using our complete set of genomes from *S. cerevisiae*, thermotolerance loci were enriched for high genotype homozygosity (Table 1 and Table S2), as seen at any given selected site after a sweep as a product of strong allele-sharing (Garud et al., 2015; Harris et al., 2018). As a control for potential demographic effects in this whole-species analysis, we repeated the test paradigm on individual well-sampled *S. cerevisiae* populations, and again detected elevated genotype homozygosity at thermotolerance genes (Tables S2 and S4). Together, our sequence-based analyses establish hallmarks of directional selection at these loci: sequence divergence between *S. cerevisiae* and *S. paradoxus*, particularly at amino-acid coding sites, and tight constraint within *S. cerevisiae*.

### Molecular evolution and functional impact of coding variation at *ESP1*

We anticipated that inspecting allele-sharing within species, at high resolution across genomic loci, could further help reveal facets of the history of thermotolerance genes. Using the largest well-sampled population of *S. cerevisiae* (collected from localities across Europe and in vineyards elsewhere), we found that genotype homozygosity was not uniform across a given thermotolerance gene, and for most loci, peaks of allele-sharing could be resolved (Figure 1A and Figure S1). As expected, the latter corresponded to troughs of polymorphism across the population as measured by the number of pairwise differences between strains (Figure S1). Notably, even at allele-sharing peaks, in absolute terms genotype homozygosity was modest. Across all thermotolerance loci, the top-scoring regions in the wine/European population was at the 5’ end of the chromosome segregation gene *ESP1* (Figure 1A), where the statistic reached at most a value of 0.15. This is consistent with our inference of an ancient date for positive selection at *ESP1* and other loci, since the tight conservation and long haplotypes expected immediately after selection would be eroded over longer timescales (Berry et al., 1991; Smith and Haigh, 1974; Weigand and Leese, 2018). Using this very highest region of allele-sharing in *ESP1* as a test case, we inspected it in other well-sampled *S. cerevisiae* populations and again found elevated genotype homozygosity (Figure 1B-E), indicating that all these populations likely have had the same forces at play at the locus.

**Figure 1.**
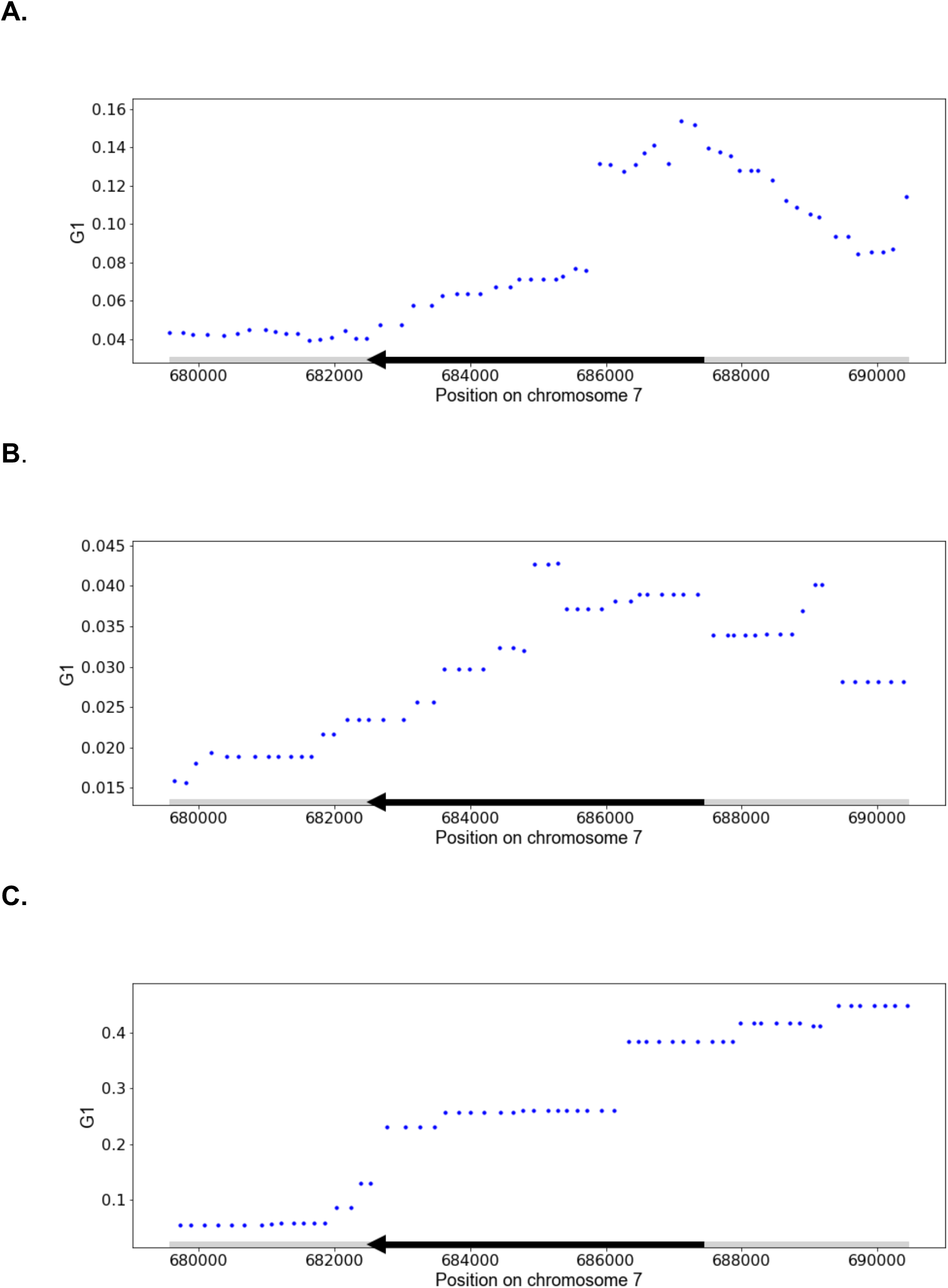
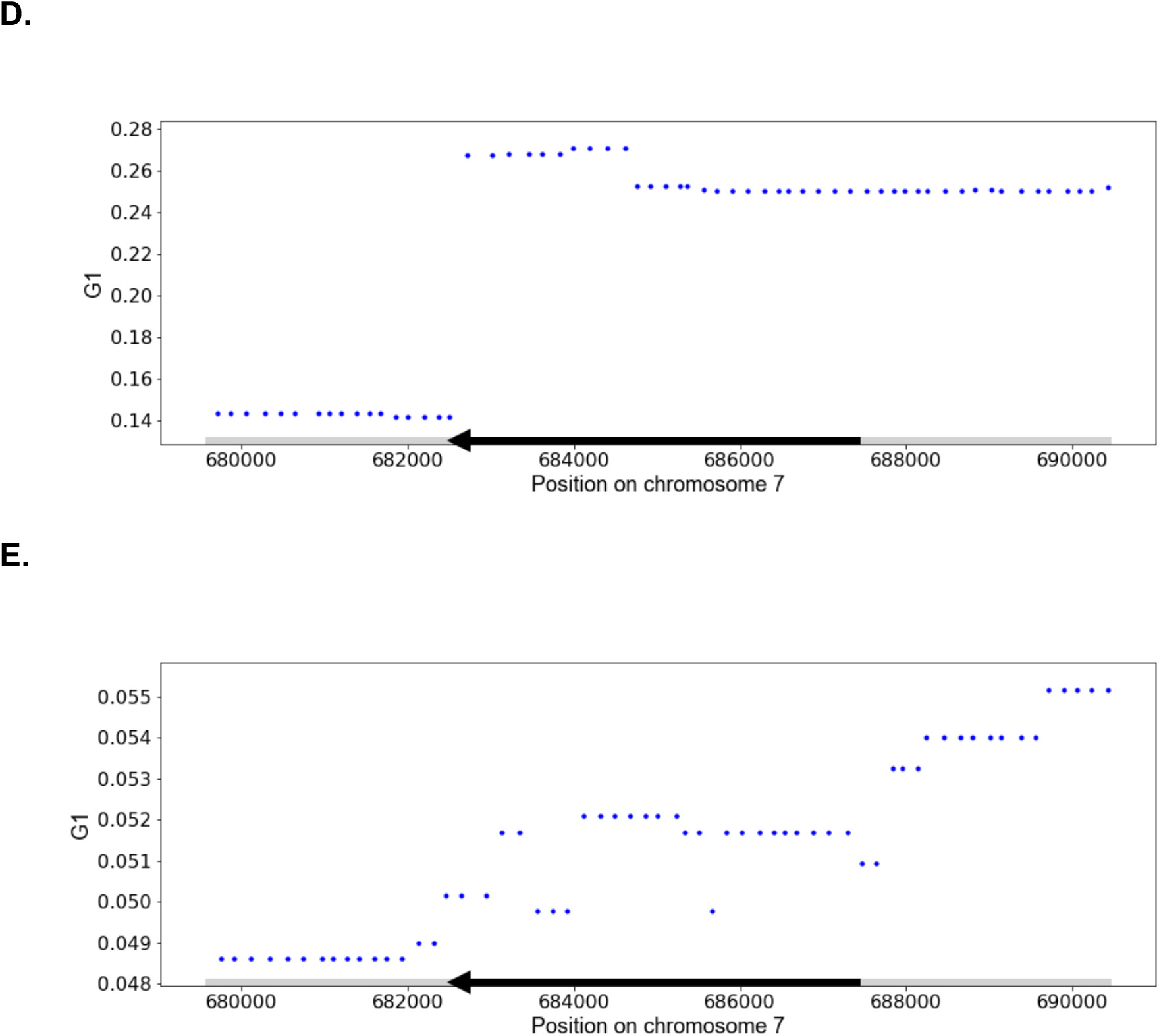
A peak of high allele frequency in *S. cerevisiae* populations at the 5’ end of *ESP1*. Each panel shows results of analysis of allele frequency at the thermotolerance gene *ESP1* in a population of *S. cerevisiae* from (Peter et al., 2018). In each panel, the *y*-axis reports genotype homozygosity, G1, in a 1200-SNP window around the position shown on the *x*. The *ESP1* open reading frame is demarcated with a dark black arrow (direction of transcription is right to left). **A**, Wine/European population. **B**, Mosaic Region 3 population. **C**, Brazilian Bioethanol population. **D**, Sake population. **E**, Mixed Origin population.

We next sought to gain deeper molecular insight into thermotolerance genetics and, for this purpose, chose to focus further on *ESP1* as a testbed, given its high allele-sharing within *S. cerevisiae* (Figure 1 and Table S2) and dramatic impact on the thermotolerance trait (Weiss et al., 2018). First, using a phylogenetic approach across the *Saccharomyces* genus, we established a pattern of accelerated protein evolutionary rate along the *S. cerevisiae* lineage in *ESP1* (*p* = 0.046), consistent with our population-level tests of protein evolution on the larger set of thermotolerance genes (Table 1). Next, to investigate the importance of coding variation at *ESP1* experimentally, we turned to an allele-swap design. We introduced the *ESP1* coding region and, separately, the *ESP1* promoter, from wild *S. paradoxus* (strain Z1, isolated from an oak tree in the United Kingdom) into a wild *S. cerevisiae* background (strain DVBPG1373, from soil in the Netherlands). Growth experiments revealed a dramatic, temperature-dependent effect of variation in the *ESP1* coding region, with the *S. paradoxus* allele compromising growth under heat treatment (Figure 2 and Figure S2). This transgenic fully recapitulated the impact of a larger, regional swap of the *S. paradoxus* open reading frame and promoter together into *S. cerevisiae* (Figure 2 and Figure S2). By contrast, the *S. paradoxus* allele of the *ESP1* promoter conferred no defect in thermotolerance when analyzed on its own (Figure 2 and Figure S2). As an independent analysis of potential promoter effects, we examined *cis*-regulatory variation between yeast species in measurements of *ESP1* gene expression. We found no overall dramatic tendency for overall *cis*-regulatory divergence between *S. cerevisiae* and other species, at *ESP1* in particular or across thermotolerance genes as a set (Table S5A). Likewise, the latter yielded no signal in tests for directional cis-regulatory divergence ((Bullard et al., 2010); Table S5B). Together, our data highlight the evolutionary and functional importance of amino acid variation between *S. cerevisiae* and *S. paradoxus* at *ESP1*, and raise the possibility that coding divergence may also underlie the thermotolerance effects of other mapped loci.

**Figure 2.**
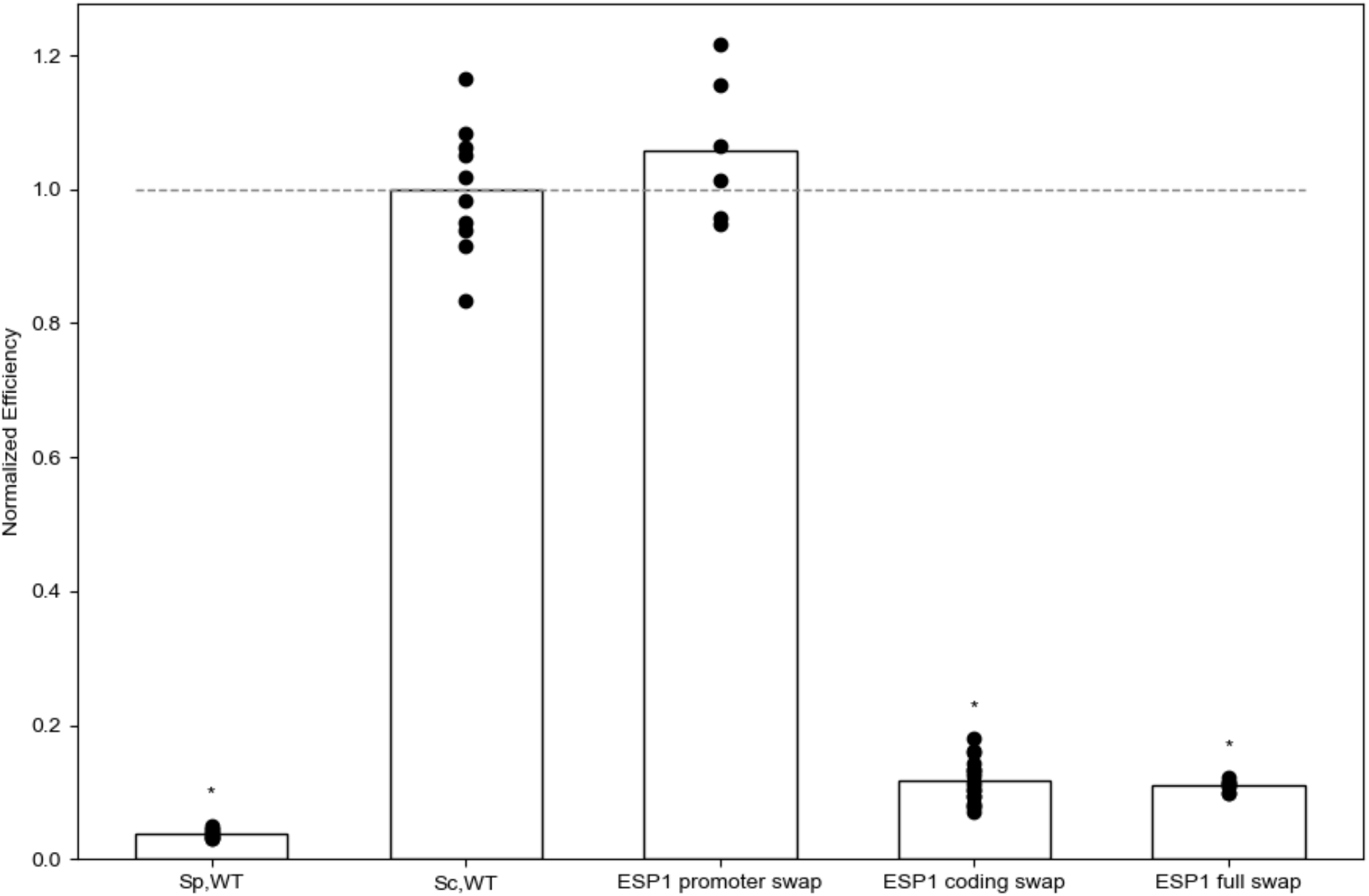
The *S. cerevisiae ESP1* coding region, but not the promoter, is required for thermotolerance. Each column represents results from biomass accumulation assays of a wild-type or transgenic yeast strain cultured at high temperature. The *y*-axis reports the optical density of a culture of the indicated strain after 24h at 39°C, normalized to the analogous quantity from wild-type *S. cerevisiae* (dashed line). Each point reports results from one biological replicate, and each bar height reports the average across replicates (*n* = 6-18). The first two columns report results from wild-type (WT) strains of *S. paradoxus* Z1 (Sp) and *S. cerevisiae* DBVPG17373 (Sc). The last three columns report strains with the indicated region of *ESP1* from *S. paradoxus* swapped into *S. cerevisiae* at the endogenous location; *ESP1* full swap denotes transgenesis of both the promoter and the coding region. *, Wilcoxon test *p* < 0.004 in a comparison against wild-type *S. cerevisiae*. Culture data at 28°C are given in Figure S2.

### Temperature dependence and cell biology of species divergence effects at *ESP1*

In further pursuit of the molecular mechanisms of *S. cerevisiae* thermotolerance, we turned to the potential for clues from temperature-dependent genetics. *S. cerevisiae* outperforms its sister species at a range of elevated temperatures (Salvadó et al., 2011; Sweeney et al., 2004). Our thermotolerance loci were identified in a screen for effects of interspecies divergence at 39°C (Weiss et al., 2018), and their relevance to growth under other conditions is unknown. Drawing again on *ESP1* as a model with which to address this question, we assayed biomass accumulation of wild-type and transgenic strains under a temperature dose-response. In these growth experiments, we observed a gradual decline in wild-type *S. cerevisiae* and *S. paradoxus* growth as temperature increased, with the latter more sensitive to heat as expected (Figure 3; AlZaben et al., 2021). Our allele-swap strain in the *S. cerevisiae* background harboring *S. paradoxus ESP1* exhibited a sharp drop in growth at ∼38°C; it grew readily below this temperature, phenocopying the wild-type *S. cerevisiae* progenitor, and at higher temperatures, it exhibited the negligible growth seen in wild-type *S. paradoxus* (Figure 3). Such a dose-response, resembling the sigmoidal behavior of a cooperative biochemical process, was a synthetic property of the *ESP1* inter-species swap, distinguishing it from either wild-type species. These data imply that, at least in the *S. cerevisiae* background, the function of *S. paradoxus* Esp1 breaks down with a steep temperature dependence, whose midpoint is close to the conditions under which this gene was originally identified (39°C).

**Figure 3.**
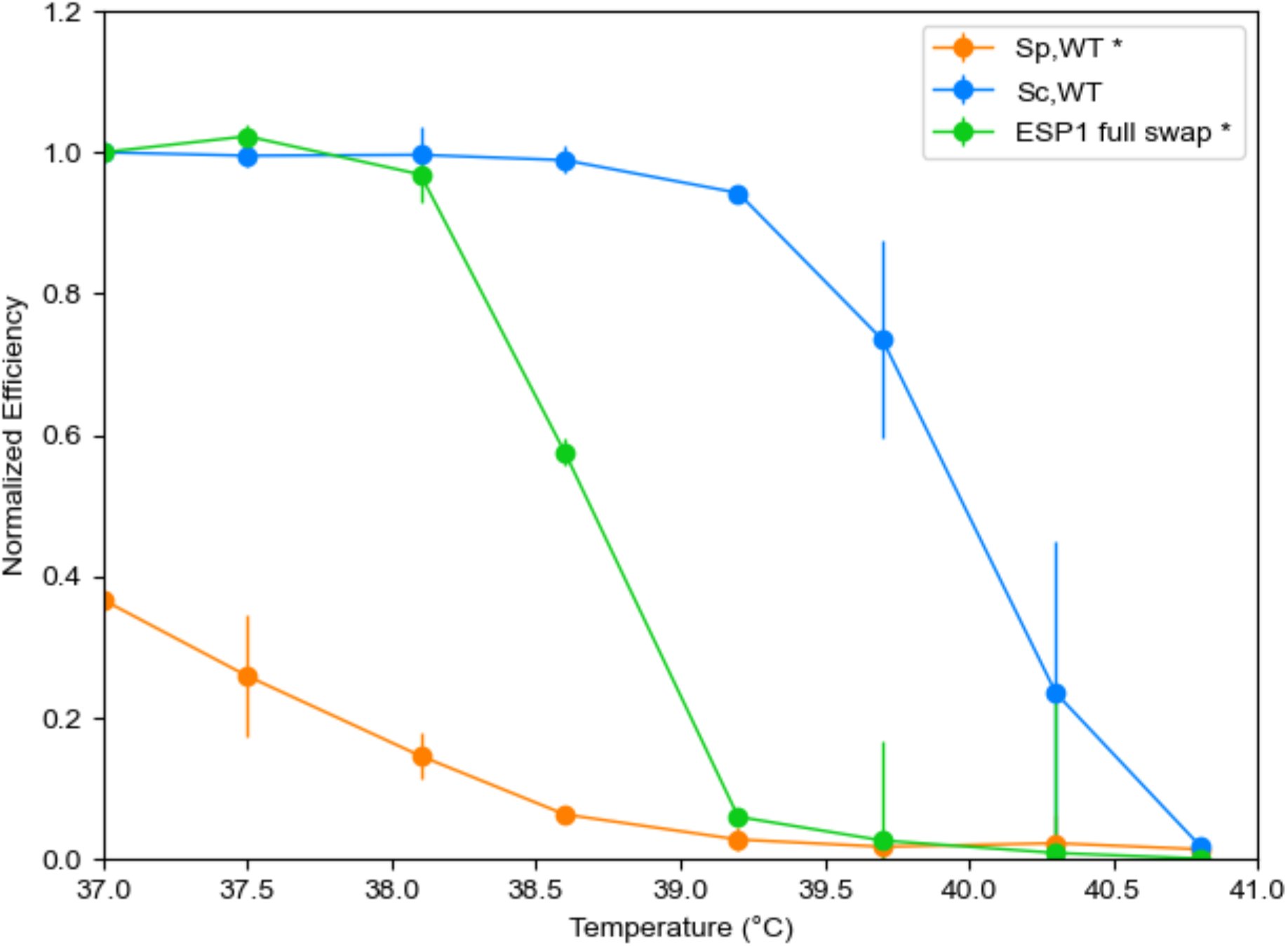
Growth function of *S. paradoxus ESP1* declines sharply with temperature. Each trace reports results from biomass accumulation assays of a wild-type or transgenic yeast strain across temperatures. Strain labels are as in Figure 2. The *y*-axis reports the optical density of a culture of the indicated strain after 24h at the temperature on the *x*, normalized to the optical density of that day’s wild-type *S. cerevisiae* at 37°C. *, *p* < 10^−12^ for the strain by temperature interaction term of a two-factor ANOVA, in a comparison between the indicated strain and wild-type *S. cerevisiae*.

*ESP1* encodes separase, which releases sister chromatids for separation into daughter cells during anaphase, cleaving the cohesin ring that has held them together in metaphase. We reasoned that, if *S. paradoxus* Esp1 failed to function, in actively growing cells harboring this allele we would see hallmarks of arrest late in the cell cycle. Quantitative microscopy bore out this prediction: as in wild-type *S. paradoxus* (Weiss et al., 2018), large-budded dyads predominated in cultures of the *S. cerevisiae* transgenic with *S. paradoxus ESP1*, when incubated at 39°C (Figure 4). These findings are consistent with a mechanism in which heat treatment compromises separase function of the *S. paradoxus* allele of Esp1, blocking the progress of the cell cycle and limiting viability and biomass accumulation. Under such a model, evolution in *S. cerevisiae* would have resolved these defects, introducing genetic changes that foster Esp1 function and boost fitness at high temperature.

**Figure 4.**
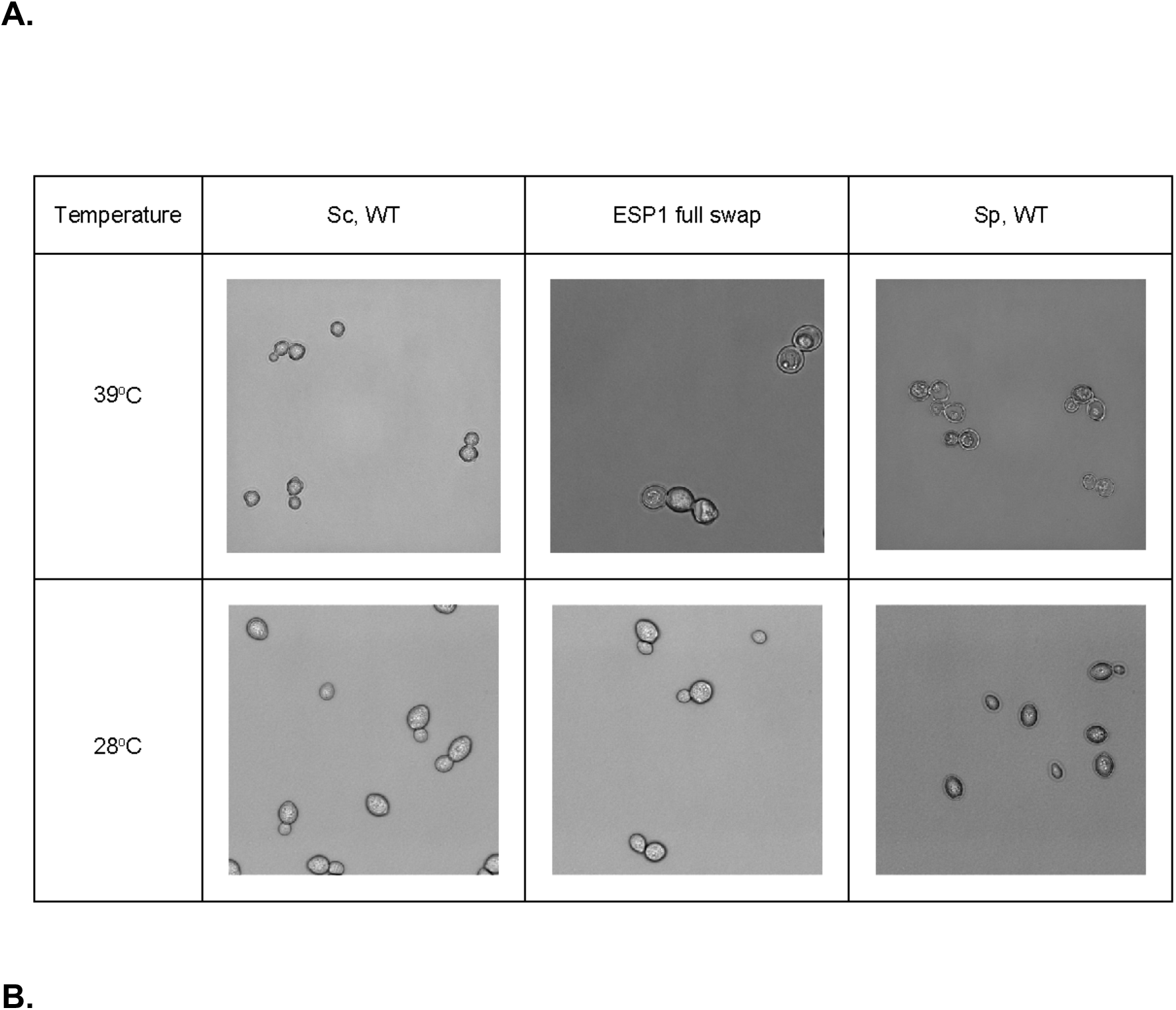
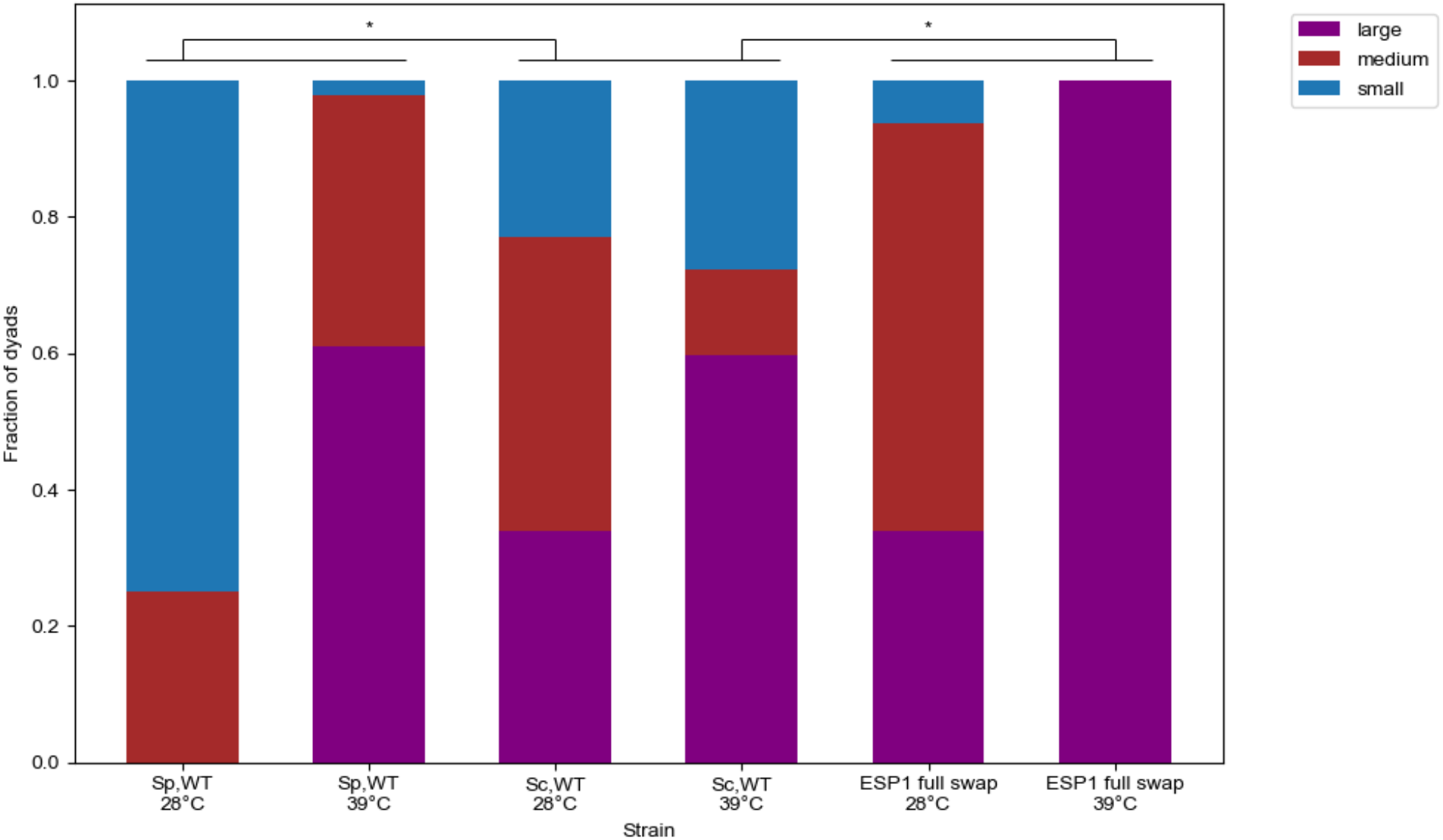
The *S. paradoxus* allele of *ESP1* compromises cell division at high temperature. **A**, Each panel reports a representative image of a wild-type or transgenic yeast strain after incubation for 24h at the indicated temperature. Strain labels are as in Figure 2. **B**, Each bar reports quantification of replicated imaging data of the indicated strain cultured at the indicated temperature, as in **A**. For each bar, the *y*-axis shows the fraction of dyads in the indicated size category. *, *p* < 0.015 for the strain by temperature interaction term of a two-factor ANOVA, in a comparison between the indicated strain and wild-type *S. cerevisiae*. Experiment details are given in Table S7.

## DISCUSSION

In the study of adaptation, a trait that arises in a species, goes to fixation, and is maintained for thousands of generations can be seen as the ultimate evolutionary success story. Here we have used yeast thermotolerance as a model of this process. We shed light on the forces driving the trait as it has evolved between species, and we investigated the molecular genetics and cell biology of divergent alleles at the underlying loci.

Our sequence-based tests of thermotolerance loci—revealing divergence and protein evolutionary rate between species, and conservation within *S. cerevisiae*—strongly suggest that the trait arose under a selective sweep before the radiation of modern *S. cerevisiae* populations. Also consistent with the latter model is the fact that at a given thermotolerance gene, alleles from *S. cerevisiae* isolates from around the world were partially sufficient for the trait, when swapped into a poorly-performing *S. paradoxus* background (Weiss et al., 2018). Plausibly, the initial rise of thermotolerance early in *S. cerevisiae* history could have been driven by the ecology of hot East Asian niches where the species likely originated (Peter et al., 2018).

The scenario of an ancient sweep of thermotolerance, whose effects bear out across modern *S. cerevisiae*, sets up an intriguing contrast with traits that undergo independent, parallel adaptations in distinct lineages of a species (Chan et al., 2010; Hoekstra and Nachman, 2003; Rosenblum et al., 2010; Xie et al., 2019). Under one compelling model, thermotolerance alleles acquired by an initially small, specialized *S. cerevisiae* ancestor could have enabled later migrants to colonize other warm niches (Robinson et al., 2016). That said, additional lineage-specific adaptations to heighten thermotolerance further could also eventually come to light.

Our data also open a window onto the molecular mechanisms of thermotolerance evolution in the yeast system. The patterns we have seen of non-synonymous sequence variation in thermotolerance genes, and our molecular-genetic experiments at the *ESP1* locus, point to a key role for protein-coding variation. What exactly would such amino acid changes be doing at thermotolerance loci? Given how much our temperature dose-response of *ESP1* function looks like a two-state protein unfolding curve, it is tempting to speculate that the *S. cerevisiae* allele of this gene may act by boosting protein stability. If such a mechanism were to prove the general rule for our loci, it would dovetail with the trend for proteome thermostability seen in heat-tolerant species (Leuenberger et al., 2017). Perhaps most likely, however, given the complexity of the trait, is a picture in which evolution tweaked many protein features (*e*.*g*. (Sas-Chen et al., 2020)), at different times and at different loci, as the *S. cerevisiae* ancestor gained its unique thermotolerance character.

## ACKNOWLEDGEMENTS AND FUNDING

The authors thank Julie Chuong for verifying and maintaining transgenic strains, David Savage for his generosity with lab facilities and resources, Josh Schraiber for discussions, and Jeremy Roop for critical reading of the manuscript. This work was supported by NSF GRFP DGE 1752814 to M.A. and NIH R01 GM120430 to R.B.B.

