## Supplemental Material for "Population and comparative genetics of thermotolerance divergence between yeast species"

| gene | p (Weiss et al, 2018)[a] | p (revised pipeline)[b] |
|---|---|---|
| **YLR397C/AFG2** | $7.0 \times 10^{-4}$ | $1.81 \times 10^{-7}$ |
| **YGR098C/ESP1** | $3.8 \times 10^{-10}$ | $4.65 \times 10^{-7}$ |
| **YMR168C/CEP3** | $5.4 \times 10^{-3}$ | $9.04 \times 10^{-6}$ |
| **YKR054C/DYN1** | $1.6 \times 10^{-6}$ | $5.16 \times 10^{-5}$ |
| **YHR023W/MYO1** | $2.8 \times 10^{-6}$ | $1.09 \times 10^{-4}$ |
| **YDR180W/SCC2** | $7.8 \times 10^{-3}$ | $7.21 \times 10^{-4}$ |
| YPL174/NIP100C | n.s. | $3.90 \times 10^{-3}$ |
| YPR164W/MMS1 | n.s. | $4.06 \times 10^{-3}$ |
| **YCR042C/TAF2** | $3.1 \times 10^{-3}$ | $4.79 \times 10^{-3}$ |
| YMR016C/SOK2 | n.s. | 0.01883 |
| YJR135C/MCM22 | n.s. | 0.02596 |
| YJL025W/RRN7 | n.s. | 0.02858 |
| YDR443C/SSN2 | n.s. | 0.03813 |
| YKL134C/OCT1 | n.s. | 0.04997 |
| # genes tested | 3416 | 4580 |
| fraction of genes tested | 0.54 | 0.72 |

**Supplementary Table 1. Re-analysis of a screen for genes at which divergence between *S. cerevisiae* and *S. paradoxus* contributes to high-temperature growth.**

[a]Published results from a reciprocal hemizygosity (RH) screen for thermotolerance in a *S. cerevisiae* x *S. paradoxus* hybrid, using an analysis scheme in which the average fitness of a given mutant across biological replicates is used as input into the RH test (Weiss et al., 2018). The first ten rows report significance of the indicated gene in the RH test (comparing thermotolerance of two sets of hemizygotes, bearing disruptions in the two species' alleles respectively, with correction for multiple testing). The bottom two rows report the number

and fraction of genes tested in this pipeline. [b]Results are as in *a* except that a distinct analysis pipeline was used, in which fitness measurements for a given mutant from biological replicates are used as independent input into the RH test. n.s., not significant at corrected $p < 0.05$.

| gene | $D_{xy}$ | NI | G1 |
|---|---|---|---|
| YLR397C/AFG2 | 0.1047 | 2.0332 | 0.0089 |
| YGR098C/ESP1 | 0.1194 | 1.9854 | 0.0157 |
| YMR168C/CEP3 | N/A | N/A | 0.0093 |
| YKR054C/DYN1 | 0.1251 | 1.9515 | 0.0147 |
| YHR023W/MYO1 | 0.1156 | 2.7887 | 0.0135 |
| YDR180W/SCC2 | 0.1215 | 1.9801 | 0.0134 |
| YPL174/NIP100C | 0.1549 | 1.4479 | 0.0190 |
| YCR042C/TAF2 | 0.1326 | 2.4492 | 0.0103 |
| YMR016C/SOK2 | 0.12 | 2.0515 | 0.0108 |
| YJR135C/MCM22 | 0.1419 | 1.8593 | 0.0064 |
| YJL025W/RRN7 | 0.1229 | 4.1086 | 0.0093 |
| YDR443C/SSN2 | 0.0992 | 2.738 | 0.0054 |
| YKL134C/OCT1 | 0.0875 | 3.9397 | 0.0056 |

**Supplementary Table 2. Metrics of interspecies diversity and intraspecies polymorphism at thermotolerance genes.**

Each row reports, for the indicated thermotolerance gene, the absolute diversity statistic $D_{xy}$, the neutrality index NI according to the McDonald-Kreitman test, and the genotype homozygosity, G1.

**A.**

| ***S. cerevisiae* population** | **Thermotolerance genes[a]** | **Genome[b]** | **p-value[c]** |
|---:|:---:|:---:|:---:|
| French Guiana Human | 0.1213 | 0.1016 | 0.0044 |
| Ale Beer | 0.1185 | 0.0996 | 0.0043 |
| West African Cocoa | 0.1203 | 0.1003 | 0.0036 |
| African Palm Wine | 0.1210 | 0.1010 | 0.0035 |
| CHNIII | 0.1206 | 0.1014 | 0.0034 |
| CHNII | 0.1199 | 0.1003 | 0.0036 |
| CHNI | 0.1214 | 0.1012 | 0.0031 |
| Taiwanese | 0.1220 | 0.1014 | 0.0020 |
| Far East Asia | 0.1201 | 0.1007 | 0.0034 |
| Malaysian | 0.1214 | 0.1012 | 0.0032 |
| Wine/European | 0.1204 | 0.1009 | 0.0030 |
| Wine/European - subclade 1 | 0.1203 | 0.1011 | 0.0037 |
| Wine/European - subclade 2 | 0.1204 | 0.1011 | 0.0039 |
| Wine/European - subclade 3 | 0.1202 | 0.1010 | 0.0037 |
| Wine/European - subclade 4 | 0.1202 | 0.1011 | 0.0040 |
| CHNV | 0.1212 | 0.1007 | 0.0027 |
| Ecuadorean | 0.1208 | 0.1007 | 0.0035 |
| Far East Russian | 0.1209 | 0.1012 | 0.0036 |
| North American Oak | 0.1206 | 0.1008 | 0.0027 |
| Asian Islands | 0.1209 | 0.1011 | 0.0030 |
| Sake | 0.1211 | 0.1012 | 0.0029 |
| Asian Fermentation | 0.1206 | 0.1010 | 0.0024 |
| Alpechin | 0.1208 | 0.1007 | 0.0035 |

| | | | |
|---|---|---|---|
| Brazilian Bioethanol | 0.1200 | 0.1003 | 0.0029 |
| Mediterranean Oak | 0.1204 | 0.1007 | 0.0038 |
| French Dairy | 0.1192 | 0.1006 | 0.0056 |
| African Beer | 0.1196 | 0.1007 | 0.0037 |
| Mosaic Beer | 0.1207 | 0.1007 | 0.0035 |
| Mixed Origin | 0.1191 | 0.0994 | 0.0037 |
| Mexican Agave | 0.1200 | 0.1007 | 0.0044 |
| M1 - Mosaic Region 1 | 0.1205 | 0.1007 | 0.0033 |
| M2 - Mosaic Region 2 | 0.1204 | 0.1007 | 0.0034 |
| M3 - Mosaic Region 3 | 0.1205 | 0.1003 | 0.0017 |

**B.**

| ***S. cerevisiae* population** | **Thermotolerance genes[a]** | **Genome[b]** | **p-value[c]** |
|---|---|---|---|
| French Guiana Human | 0.1259 | 0.1032 | 0.0016 |
| Ale Beer | 0.1226 | 0.1015 | 0.0035 |
| West African Cocoa | 0.1258 | 0.1021 | 0.0009 |
| African Palm Wine | 0.1266 | 0.1026 | 0.0019 |
| CHNIII | 0.1253 | 0.1031 | 0.0018 |
| CHNII | 0.1249 | 0.1020 | 0.0014 |
| CHNI | 0.1271 | 0.1026 | 0.0007 |
| Taiwanese | 0.1271 | 0.1031 | 0.0010 |
| Far East Asia | 0.1243 | 0.1024 | 0.0014 |
| Malaysian | 0.1271 | 0.1028 | 0.0008 |
| Wine/European | 0.1228 | 0.1027 | 0.0056 |

| | | | |
|---|---|---|---|
| Wine/European - subclade 1 | 0.1252 | 0.1027 | 0.0022 |
| Wine/European - subclade 2 | 0.1252 | 0.1029 | 0.0018 |
| Wine/European - subclade 3 | 0.1252 | 0.1028 | 0.0014 |
| Wine/European - subclade 4 | 0.1251 | 0.1028 | 0.0025 |
| CHNV | 0.1264 | 0.1028 | 0.0016 |
| Ecuadorean | 0.1266 | 0.1025 | 0.0003 |
| Far East Russian | 0.1278 | 0.1028 | 0.0009 |
| North American Oak | 0.1265 | 0.1025 | 0.0009 |
| Asian Islands | 0.1266 | 0.1027 | 0.0008 |
| Sake | 0.1278 | 0.1028 | 0.0007 |
| Asian Fermentation | 0.1261 | 0.1026 | 0.0007 |
| Alpechin | 0.1259 | 0.1026 | 0.0012 |
| Brazilian Bioethanol | 0.1251 | 0.1021 | 0.0009 |
| Mediterranean Oak | 0.1254 | 0.1025 | 0.0014 |
| French Dairy | 0.1236 | 0.1024 | 0.0015 |
| African Beer | 0.1243 | 0.1026 | 0.0017 |
| Mosaic Beer | 0.1257 | 0.1026 | 0.0006 |
| Mixed Origin | 0.1239 | 0.1013 | 0.0017 |
| Mexican Agave | 0.1249 | 0.1023 | 0.0016 |
| M1 - Mosaic Region 1 | 0.1256 | 0.1026 | 0.0007 |
| M2 - Mosaic Region 2 | 0.1255 | 0.1025 | 0.0023 |
| M3 - Mosaic Region 3 | 0.1259 | 0.1020 | 0.0011 |

**Supplementary Table 3. Divergence between *S. cerevisiae* and *S. paradoxus* at thermotolerance genes.**

**A.** [a]Median absolute divergence ($D_{xy}$) between the indicated population of *S. cerevisiae* from (Peter et al., 2018) and the Wine/European *S. paradoxus* population from (Bergström et al., 2014) in thermotolerance genes. [b]Genomic median $D_{xy}$ for each population pair as in *a.* [c]Empirical significance from a resampling test.

**B.** Data are as in **A**, except that each comparison is between the indicated population of *S. cerevisiae* from (Peter et al., 2018) and the North American *S. paradoxus* subpopulation B from (Durand et al., 2019).

| Population | # Isolates | Thermotolerance genes[a] | Genome[b] | *p*-value[c] |
|---|---|---|---|---|
| Wine/European | 362 | 0.038 | 0.026 | 0.0526 |
| Mosaic Region 3 | 113 | 0.020 | 0.019 | 0.200 |
| Mixed Origin | 72 | 0.076 | 0.061 | 0.0493* |
| Sake | 47 | 0.265 | 0.258 | 0.280 |
| Brazilian Bioethanol | 35 | 0.131 | 0.109 | 0.0253* |

**Supplementary Table 4. Thermotolerance loci are enriched for allele-sharing within *S. cerevisiae* populations.**

Each row reports results from analyses of the allele-sharing statistic genotype homozygosity, G1, for the indicated population of *S. cerevisiae* from (Peter et al., 2018). [a]Median G1 in thermotolerance genes. [b]Genomic median G1. [c]Empirical significance from a resampling test. *, $p < 0.05$.

**A**

| | Genome | Thermotolerance loci | *p*-value |
|---|---|---|---|
| ***S. paradoxus***[a] | 0.197 | 0.138 | 0.8315 |
| ***S. uvarum***[b] | 0.073 | 0.089 | 0.6912 |

**B**

| | Genome | Thermotolerance loci | *p*-value |
|---|---|---|---|
| ***S. paradoxus***[a] | -0.011 | 0.138 | 0.2086 |
| ***S. uvarum***[b] | 0.001 | -0.029 | 0.4362 |

**Supplementary Table 5. No consistent *cis*-regulatory divergence between *S. cerevisiae* and other species at thermotolerance loci.**

**A.** [a]Median *cis*-regulatory divergence in thermotolerance genes between alleles of the indicated species in their F1 hybrid. [b]Data are as in [a] except that the value reports an average across random sets of genes from the genome. [c]Results of a one-sided resampling test for an elevated magnitude of *cis*-regulatory divergence in thermotolerance genes.

**B.** Data are as in **A** except that the test was for directional *cis*-regulatory divergence across thermotolerance genes.

| Species background | Strain background | Swap (if applicable)[a] | Swap boundaries[a] | Source | Strain Name |
|---|---|---|---|---|---|
| *S. paradoxus* x *S. cerevisiae* | Z1 x DBVPG1373 | N/A | N/A | Weiss et al., 2018 | CW27 |
| *S. paradoxus* | Z1 | N/A | N/A | Weiss et al., 2018 | CW62 |
| *S. cerevisiae* | DBVPG1373 | N/A | N/A | Weiss et al., 2018 | CW68 |
| *S. cerevisiae* | DBVPG1373 | ESP1 full swap | Chr7:687097-682525 | Weiss et al., 2018 | CW98 |
| *S. cerevisiae* | DBVPG1373 | ESP1 promoter | Chr7:687463-687842 | This study | CW339 |
| *S. cerevisiae* | DBVPG1373 | ESP1 coding swap | Chr7:682525-687419 (heterozygous at position 687419) | This study | CW412 |
| *S. cerevisiae* | DBVPG1373 | ESP1 coding swap | Chr7:682525-687419 | This study | CW413 |
| *S. cerevisiae* | DBVPG1373 | ESP1 coding swap | Chr7:682525-687419 | This study | CW414 |

**Supplementary Table 6. Strains used in this study**.

[a]For strains harboring *ESP1* from *S. paradoxus* swapped into the *S. cerevisiae* background at the endogenous location, listed is the mode of transgenesis (where full indicates both the promoter and coding regions) and boundaries of swapped genetic material. N/A, not applicable.

| Strain | Temperature | Minimum[a] | Maximum[a] | Median[a] | Total[b] | Images[c] |
|---|---|---|---|---|---|---|
| *Sp,* WT | 28°C | 0 | 2 | 0 | 5 | 21 |
| | 39°C | 0 | 6 | 2 | 42 | 21 |
| *Sc*, WT | 28°C | 0 | 4 | 2 | 37 | 19 |
| | 39°C | 0 | 3 | 1 | 15 | 16 |
| ESP1 full swap | 28°C | 0 | 6 | 2 | 32 | 19 |
| | 39°C | 0 | 2 | 0 | 13 | 27 |

**Supplementary Table 7. Details of microscopy experiments analyzing cell division in wild-type and *ESP1* transgenic yeast.**

Each row reports the details, from Figure 4 of the main text, of analyses of microscopy of the indicated strain cultured at the indicated temperature. [a]Minimum, maximum, and median dyad count per image. [b]Total number of dyads counted across images. [c]Total number of images scored.

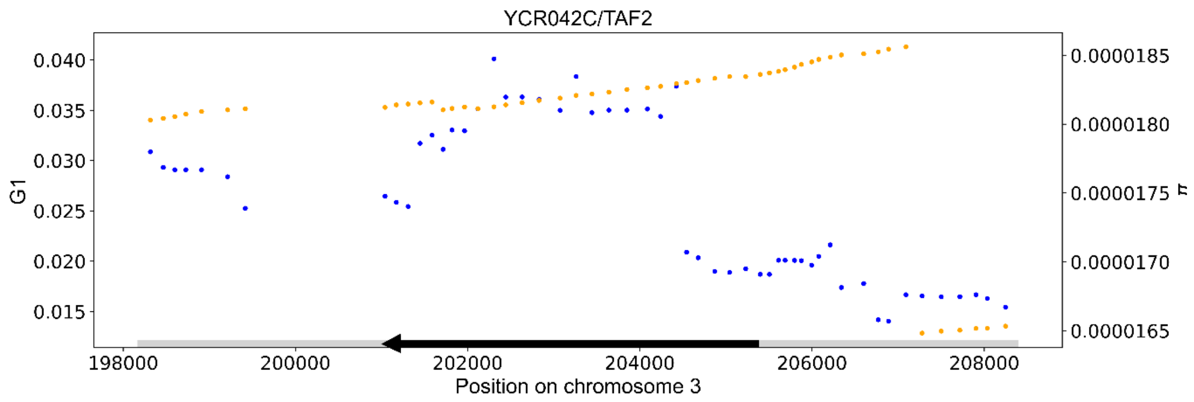
YCR042C/TAF2
G1
π
Position on chromosome 3

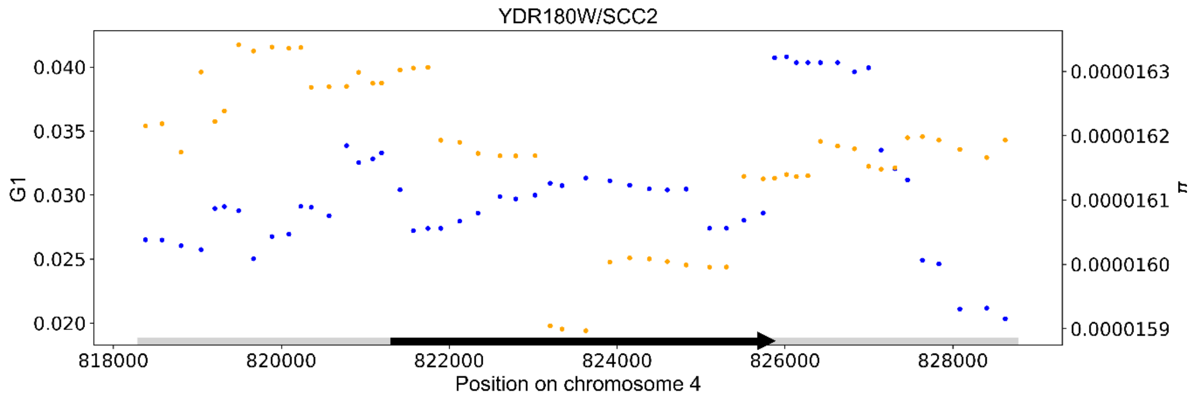
YDR180W/SCC2
G1
π
Position on chromosome 4

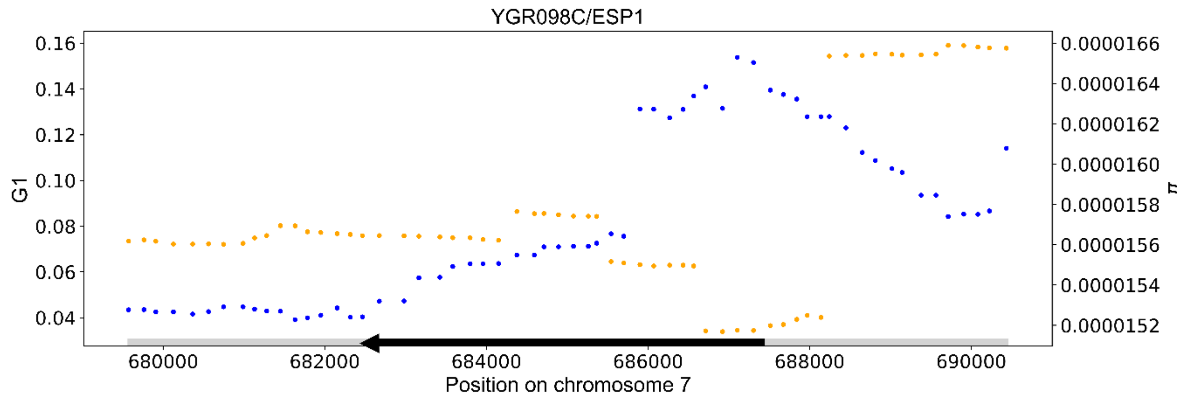
YGR098C/ESP1
G1
π
Position on chromosome 7

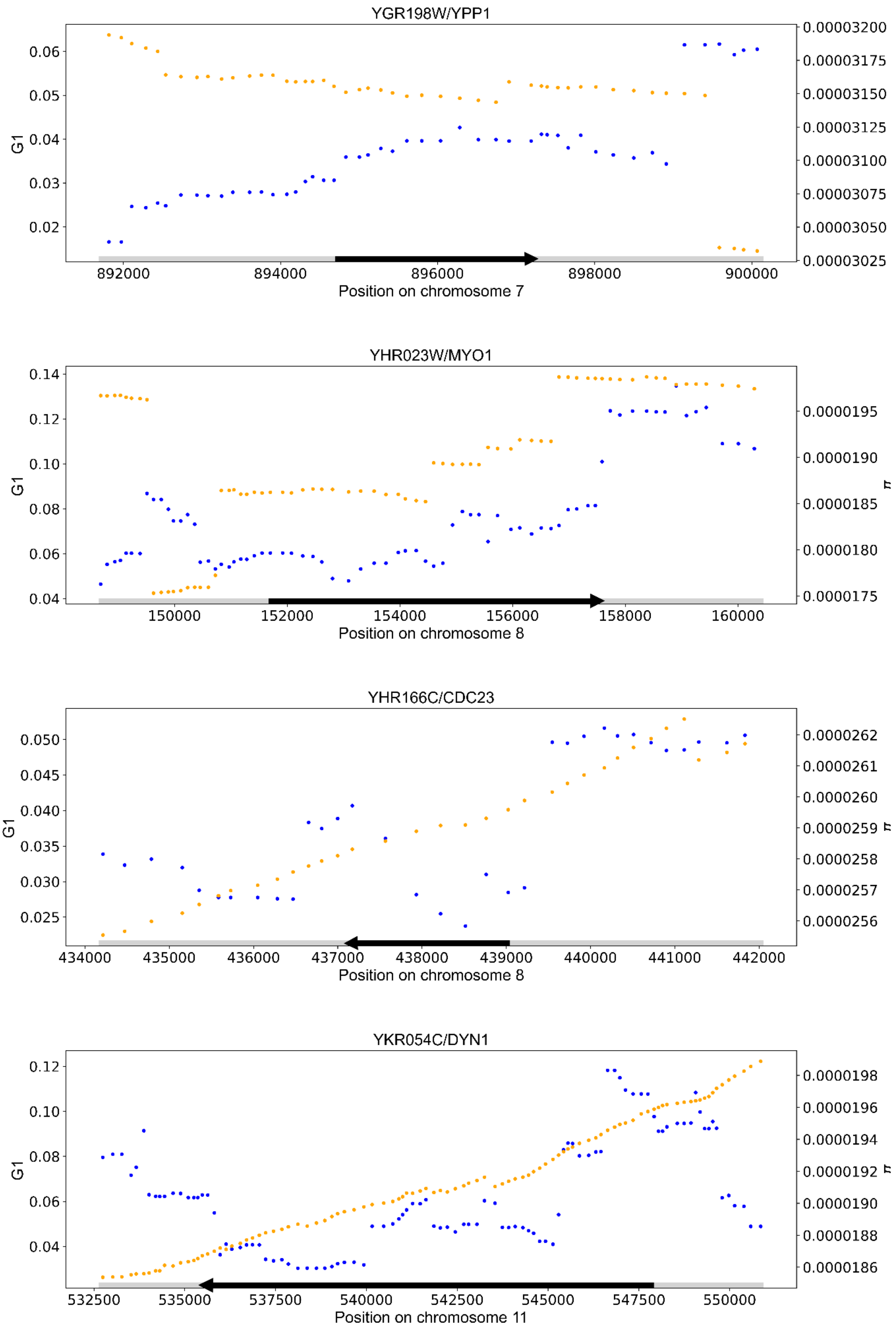
YGR198W/YPP1
G1
0.06
0.05
0.04
0.03
0.02
0.00003200
0.00003175
0.00003150
0.00003125
0.00003100
0.00003075
0.00003050
0.00003025
892000
894000
896000
898000
900000
Position on chromosome 7
YHR023W/MYO1
G1
0.14
0.12
0.10
0.08
0.06
0.04
0.0000195
0.0000190
0.0000185
0.0000180
0.0000175
π
150000
152000
154000
156000
158000
160000
Position on chromosome 8
YHR166C/CDC23
G1
0.050
0.045
0.040
0.035
0.030
0.025
0.0000262
0.0000261
0.0000260
0.0000259
0.0000258
0.0000257
0.0000256
π
434000
435000
436000
437000
438000
439000
440000
441000
442000
Position on chromosome 8
YKR054C/DYN1
G1
0.12
0.10
0.08
0.06
0.04
0.0000198
0.0000196
0.0000194
0.0000192
0.0000190
0.0000188
0.0000186
π
532500
535000
537500
540000
542500
545000
547500
550000
Position on chromosome 11

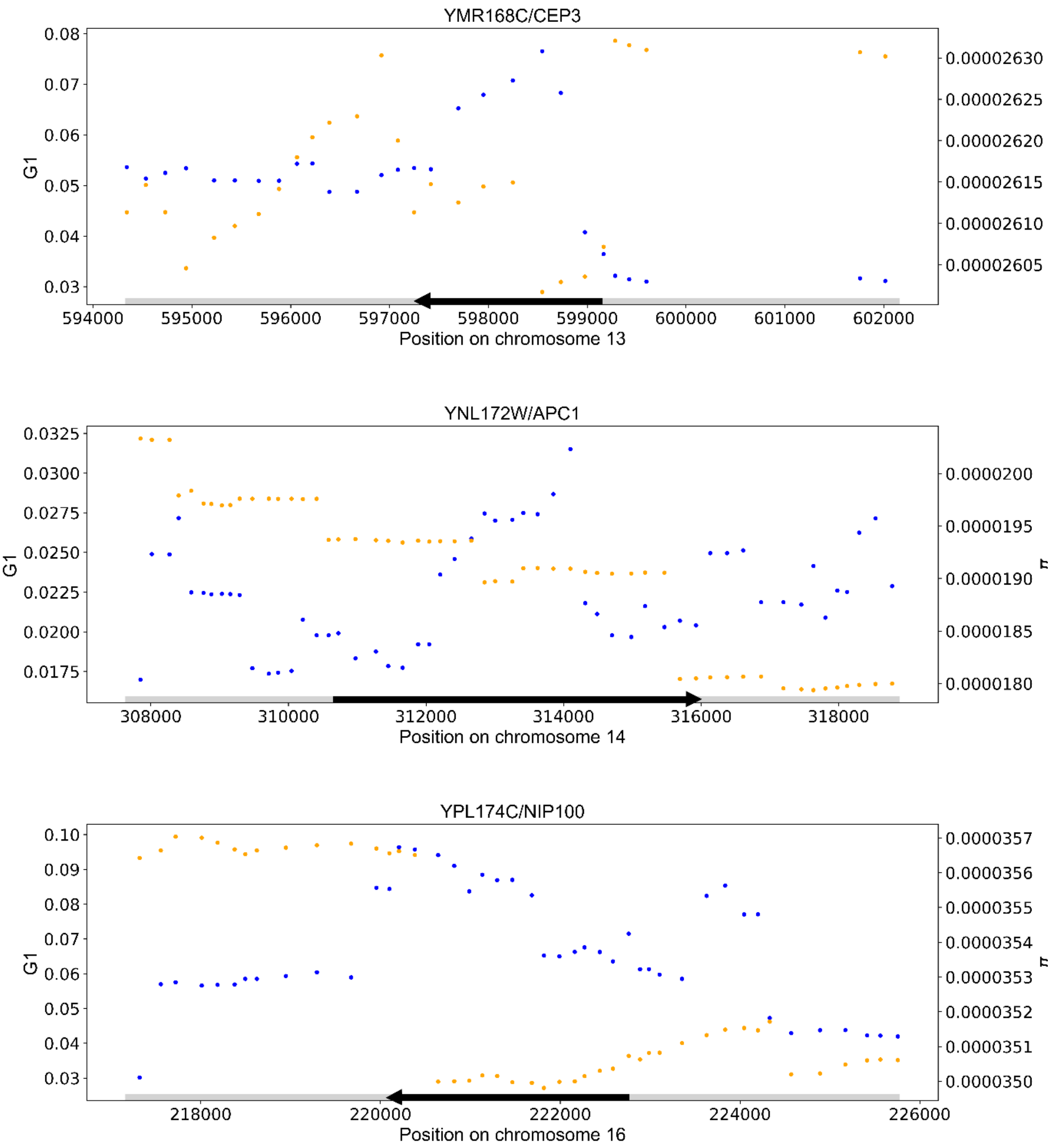


**Supplementary Figure 1. Spatial distribution of allele sharing in wine/European *S. cerevisiae* at thermotolerance genes.** Data are as in Figure 1A of the main text, except that each panel reports results for one thermotolerance gene, and the right *y*-axis reports nucleotide diversity (π, orange dots) in each 1200-SNP window centered around the position on the *x*.

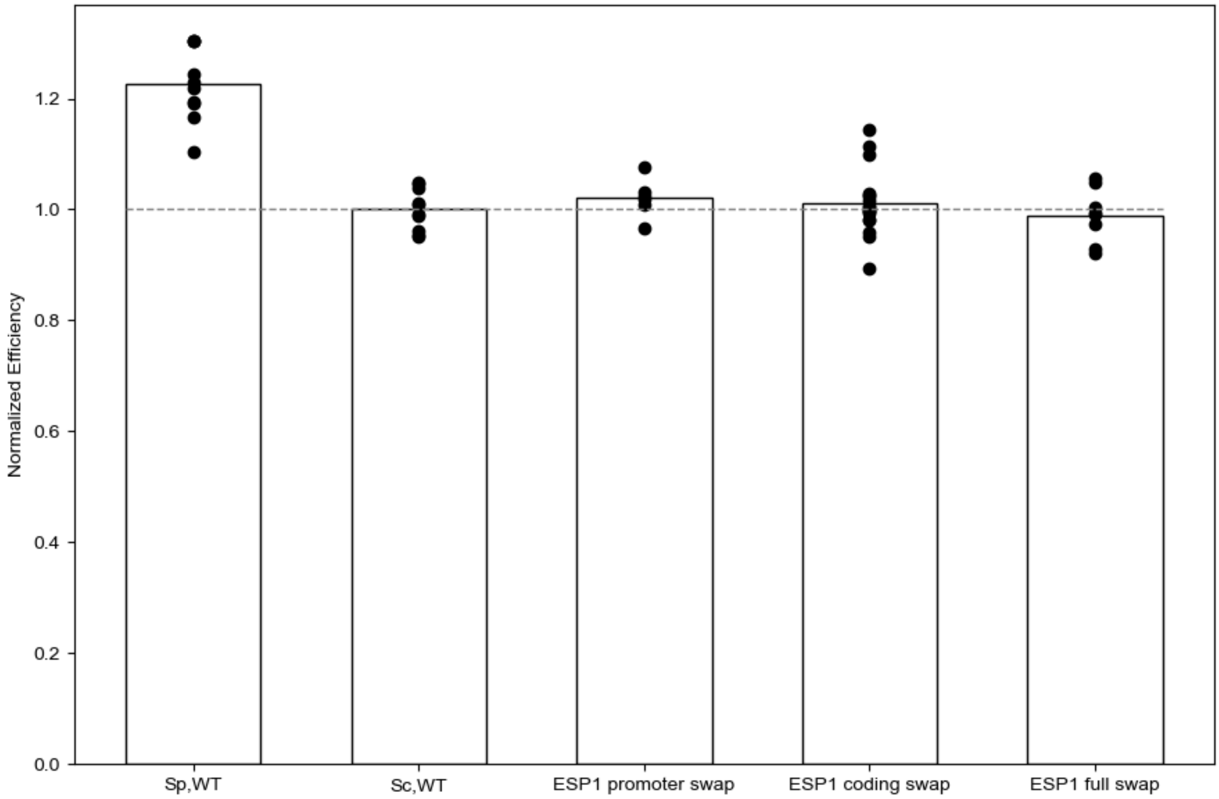


**Supplementary Figure 2. Strain background has no impact on growth at 28°C.** Data are as in Figure 2 of the main text, except that culture experiments were done at 28°C (*n* = 6-18). No strains exhibited a significant growth decrease compared to wild-type *S. cerevisiae*.
